# Neural excitation/inhibition imbalance and the treatment of severe depression

**DOI:** 10.1101/2021.07.09.451784

**Authors:** Freek ten Doesschate, Willem Bruin, Peter Zeidman, Christopher C. Abbott, Miklos Argyelan, Annemieke Dols, Louise Emsell, Philip F.P. van Eijndhoven, Eric van Exel, Peter C.R. Mulders, Katherine Narr, Indira Tendolkar, Didi Rhebergen, Pascal Sienaert, Mathieu Vandenbulcke, Joey Verdijk, Mike van Verseveld, Hauke Bartsch, Leif Oltedal, Jeroen A. van Waarde, Guido A. van Wingen

**Affiliations:** Department of Psychiatry, Rijnstate hospital, Arnhem, the Netherlands; Amsterdam UMC, University of Amsterdam, Department of Psychiatry, Amsterdam Neuroscience, Amsterdam, the Netherlands; Wellcome Centre for Human Neuroimaging, 12 Queen Square, London, WC1N 3AR, UK; Department of Psychiatry, University of New Mexico School of Medicine, Albuquerque, New Mexico; Center for Psychiatric Neuroscience at the Feinstein Institute for Medical Research, New York, New York; GGZ inGeest Specialized Mental Health Care, Department of Old Age Psychiatry, Oldenaller 1, 1081 HJ Amsterdam, the Netherlands; Amsterdam UMC, Vrije Universiteit, Psychiatry, Amsterdam Public Health Research Institute, the Netherlands; Amsterdam UMC, Vrije Universiteit, Psychiatry, Amsterdam Neuroscience, the Netherlands; Katholieke Universiteit Leuven, University Psychiatric Center Katholieke Universiteit Leuven, Leuven, Belgium; Donders Institute for Brain, Cognition and Behavior, Department of Psychiatry, Nijmegen, the Netherlands; Department of Psychiatry, Radboud University Medical Centre, Huispost 961, P.O. Box 9101, 6500 HB Nijmegen, the Netherlands; Departments of Neurology, Psychiatry, and Biobehavioral Sciences, University of California, Los Angeles, California; Academic Center for ECT and Neuromodulation (AcCENT), University Psychiatric Center KU Leuven (Catholic University of Leuven), Leuven, Belgium; Mohn Medical Imaging and Visualization Center, Department of Radiology, Haukeland University Hospital, Bergen, Norway; Department of Clinical Medicine, University of Bergen, Bergen, Norway; Center for Multimodal Imaging and Genetics, University of California, San Diego, La Jolla, California; Department of Radiology, University of California, San Diego, La Jolla, California; Amsterdam Brain and Cognition, University of Amsterdam, the Netherlands

## Abstract

An influential hypothesis holds that depression is related to a neural excitation/inhibition imbalance, but its role in the treatment of depression remains unclear. Here, we show that unmedicated patients with severe depression demonstrated reduced inhibition of brain-wide resting-state networks relative to healthy controls. Patients using antidepressants showed inhibition that was higher than unmedicated patients and comparable to controls, but they still suffered from severe depression. Subsequent treatment with electroconvulsive therapy (ECT) reduced depressive symptoms, but its effectiveness did not depend on changes in network inhibition. Concomitant pharmacotherapy increased the effectiveness of ECT, but only when the strength of neural inhibition before ECT was within the normal range and not when inhibition was excessive. These findings suggest that reversing the excitation/inhibition imbalance may not be sufficient nor necessary for the effective treatment of severe depression, and that brain-state informed pharmacotherapy management may enhance the effectiveness of ECT.

## Main

With more than 250 million people affected, depression is one of the leading causes of disability worldwide^1^. Patients suffer weeks to sometimes years of low mood, anhedonia, sleep problems, weight loss, and — in more severe cases — suicidality, motor retardation and psychotic features. Although 70% of patients show a positive response to (extensive) treatments with pharmacotherapy and psychotherapy^2^, approximately half of them eventually relapse and experience one or more recurrent episodes in their lifetime^2,3^. Patients suffering from severe depressive episodes that are not responsive to initial treatments may benefit from electroconvulsive therapy (ECT). With a remission rate of 48-65%, ECT is currently the most effective treatment for pharmacotherapy-resistant depression^4^. Nevertheless, it is clear that for a substantial group of depressed patients an effective treatment is still lacking. Enhancing our understanding on the neural mechanisms of depression and its treatments is indispensable to improve treatment effectiveness and to develop novel treatments.

Healthy functioning of cortical brain circuits depends on processes that promote homeostasis throughout the network, which is regulated by excitatory and inhibitory neurotransmission^5^. One of the proposed neural mechanisms of depression states that symptoms are caused by deficits in inhibitory neurotransmission, resulting in an excitation/inhibition imbalance^6–9^. In line with this hypothesis, deficits in the principal inhibitory neurotransmitter γ-aminobutyric acid (GABA) seem to cause depression-like behavior in animal models^10,11^. Furthermore, GABA concentrations are reduced in depressed patients but not in remitted patients^12^. Accordingly, this hypothesis also predicts that reversing the excitation/inhibition imbalance in depressed patients will lead to a reduction of depressive symptoms^7,9,10^. Indeed, even though classical antidepressants primarily target other neurotransmitters, these agents seem to stimulate GABA-ergic neurotransmission as well^6,7,9,13,14^. Furthermore, benzodiazepines (i.e., a GABA_A_ receptor agonist) provide early antidepressant effects when combined with antidepressants^15^. Thus, this hypothesis poses that the neural excitation/inhibition imbalance takes a central position in the pathophysiology of depression and its treatment.

Similar to pharmacotherapy, the effectiveness of ECT may also depend on mechanisms that are closely related to the neural excitation/inhibition balance^16^. ECT uses a brief electrical stimulus to the cranium in order to elicit seizure activity which propagates throughout the brain^17^. For a beneficial treatment response, the administered stimulus should significantly exceed the individuals’ seizure threshold^18^, thereby providing strong excitation of the brain. Furthermore, ECT effectiveness depends on the generalization of the induced seizure^19^, which is accomplished through the opposing forces of excitatory seizure activity and inhibitory restraints^20^. Also, ECT effectiveness is associated with the strength of the post-ictal suppression of induced seizure activity^21^. This phenomenon is marked by strong inhibitory processes terminating the excitatory seizure dynamics. Thus, the working mechanisms of ECT seem to be tightly linked to the excitation/inhibition balance, indicating that disturbances in this balance may influence ECT effectiveness.

In order to investigate the excitation/inhibition balance in depression and its relation to treatment effects, we used data from a large multicenter international consortium that included patients with severe depression who were treated with pharmacotherapy and subsequent ECT. We used spectral dynamic causal modeling (spDCM) of resting-state functional magnetic resonance imaging (RS-fMRI) data^22^. spDCM is able to accurately estimate extrinsic (i.e., between-region) and intrinsic (i.e., regional self-inhibition) effective connectivity (Figure 1)^22–24^. Well-tuned regional self-inhibition allows for sustained activity in the cortex while maintaining a balance between excitation and inhibition throughout the network. The strength of intrinsic self-inhibition can thus be interpreted as a proxy for the regional excitation/inhibition balance^25^. Recently, a study using magnetoencephalography showed that DCM can quantify the increase in tonic inhibition by a GABA reuptake inhibitor^26^. Here, we use spDCM to study the effects of pharmacotherapy and ECT on extrinsic and intrinsic effective connectivity in severely depressed patients. We provide evidence indicating that reversing the neural excitation/inhibition imbalance is, in and of itself, not sufficient nor necessary for an antidepressant response in patients suffering from severe depression. Our findings also suggest that concomitant pharmacotherapy increases the effectiveness of ECT if the strength of inhibition remains within a healthy range, but not when the degree of inhibition is excessive.

**Figure 1.**
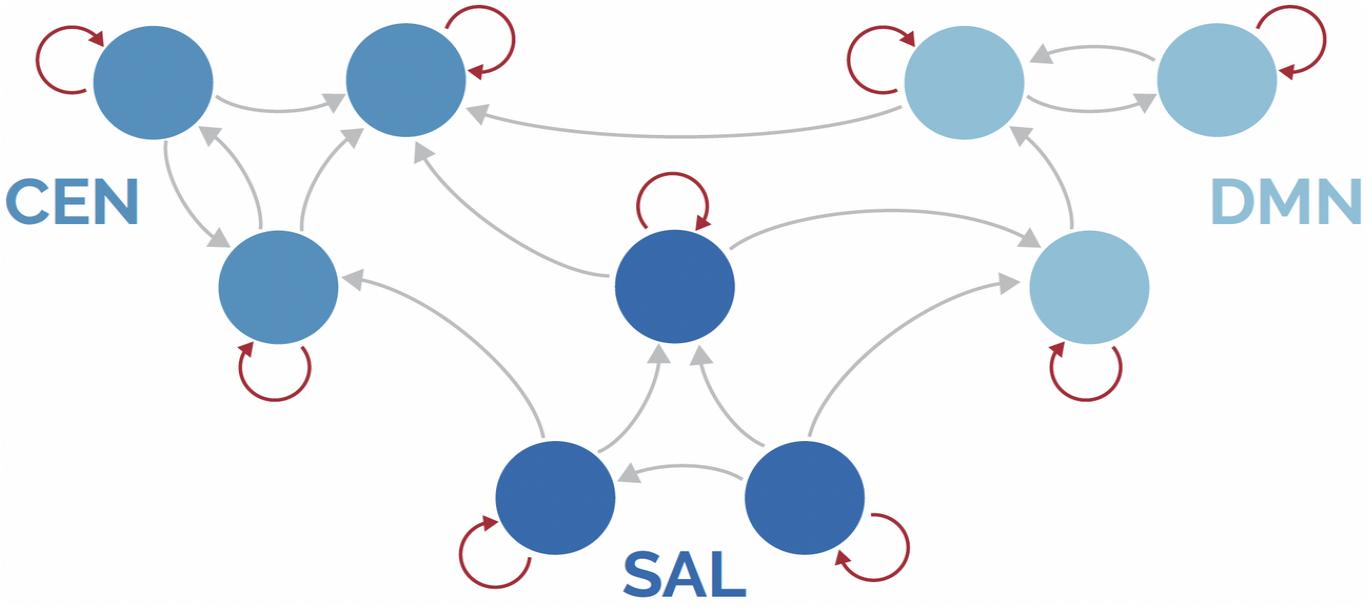
Schematic representation of a directed cortical network. The nodes (blue circles) represent cortical regions of brain networks that interact through effective connections (gray arrows). The self-connections (red arrows) regulating the strength of regional inhibition. Optimal strength of self-connections allows for sustained activity in the cortex while maintaining a balance between excitation and inhibition throughout the cortical network. The selected networks in this study are the central executive network (CEN), salience network (SAL), and default mode network (DMN).

## Results

### Reversal of the excitation/inhibition imbalance is not sufficient for pharmacotherapy effectiveness in severe depression

Data were obtained from the Global ECT-MRI Research Collaboration (GEMRIC) database^27^. Participants from seven sites across Europe and North America were divided into four groups: healthy controls (n = 59), unipolar depressed patients without current use of antidepressants or benzodiazepines (patients might use other types of medication, but for convenience were labeled as ‘unmedicated group’; n = 60), unipolar depressed patients using antidepressants (n = 56), and unipolar depressed patients using benzodiazepines (n = 63). The latter two groups included patients using both types of medication (n = 24), this considerable overlap was accounted for in the analysis (see SI Methods). Demographic and clinical characteristics of each group are reported in Table 1.

**Table 1.** Overview of demographic and clinical information. Mean (±SD) and count (%) statistics of variables for each specific group. Analyses on group differences were performed using student’s t-test or X^2^-test, where appropriate. HC = healthy controls; AD = patients using antidepressants; BZ = patients using benzodiazepines; NM = patients without current AD or BZ medication use; HAM-D = Hamilton Rating Scale for depression; ^a^P_HC vs AD_ < 0.1; ^b^P_HC vs NM_ < 0.1; ^c^P_HC vs BZ_ < 0.1; ^d^P_NM vs AD_ < 0.1; ^e^P_NM vs BZ_ < 0.1; ^f^P_AD vs BZ_ < 0.1.

|  |  |  |  | Overlap = 24 |  |
| --- | --- | --- | --- | --- | --- |
|  | Statistic | HC (n = 59) | NM (n = 60) | AD (n = 56) | BZ (n = 63) |
| **Age** <sup>a,c,d,e,f</sup> | Mean (+- 1SD) | 48.1 (15.1) | 48.7 (16.1) | 63.1 (11.5) | 54.9 (14.3) |
| **Sex** | Count (%) | 26 (44.1) | 25 (41.7) | 19 (33.9) | 29 (46) |
| **HAM-D** <sup>e</sup> | Mean (+- 1SD) | NA | 23.8 (6.1) | 25.2 (6.6) | 27.6 (7.1) |
| **Antipsychotics use** <sup>d,e</sup> | Count (%) | NA | 4 (6.7) | 29 (51.8) | 25 (39.7) |
| **Lithium carbonate use** | Count (%) | NA | 1 (1.7) | 4 (7.1) | 2 (3.2) |
| **Recurrent depression** | Count (%) | NA | 56 (93.3) | 53 (94.6) | 59 (93.7) |
| **Psychotic features** <sup>d,e</sup> | Count (%) | NA | 8 (13.3) | 17 (30.4) | 10 (15.9) |

We investigated whether these groups differed in effective connectivity using RS-fMRI and spDCM (see SI Methods)^22^. On the individual level, spDCM modeled causal interactions between neuronal populations located in the regions of interest (ROIs). A forward model generated simulated RS-fMRI timeseries based on these neural models. The cross-spectra of these simulated timeseries were compared to cross-spectra of observed RS-fMRI timeseries. Specifically, Bayesian statistical methods identified neural connectivity parameters that maximized model accuracy (the similarity of the simulated and observed cross-spectra), while minimizing complexity (the effective number of parameters). Using parametric empirical Bayes, the connectivity parameters were then taken to the group level and modeled using a general linear model (GLM; for model designs, see SI Methods and SI Figures). Thereby, spDCM enabled inference on effective (i.e., directed) connectivity between brain regions (Figure 1). Additionally, it allowed inference on the strength of regional inhibitory self-connections, which could be interpreted as local regulation of the excitation/inhibition ratio in order to maintain balance throughout the network (Figure 1)^25^. Fourteen ROIs were selected, located at the core of the default mode network (DMN), salience network (SN) and central executive network (CEN; see SI Figure 1). These networks are implicated in the pathophysiology of depression^28^. The analyses yielded no evidence for effects in between-region connectivity, so we focused on the effects in self-connections below. By comparing the evidence for the full connectivity GLM against specific reduced GLMs (SI Methods), we examined effects in inhibitory selfconnections of individual ROIs. Also, we examined effects at the network-level, which tested if effects on self-inhibition were present in none of the networks (i.e., evidence for absence of an effect), in a specific network, or widespread across networks (see SI Figure 3 and SI Methods). The degree of evidence was expressed as the probability of an effect in a (combination of) connection(s) given the data (i.e., the posterior probability, written as p(m|Y)). Evidence was considered strong if the posterior probability was 95% or higher (i.e., p(m|Y) > 0.95).

Because multiple GLM designs could be used to examine cross-sectional group differences, we tested the two most convenient designs and selected the one which maximized model evidence (SI Figure 2). The winning model design consisted of separate regressors for each patient group, while healthy controls were modeled as the reference group (i.e., intercept). The results of the selected model showed that self-inhibition was reduced across networks in unmedicated patients compared to healthy controls (p(m_4_|Y) = 1.0). Patients using benzodiazepines showed an opposite pattern of increased selfinhibition across networks compared to healthy controls (p(m_4_|Y) = 1.0). Using Bayesian statistics allowed us to establish the evidence for a null finding. In this case, the results suggest that self-inhibition in patients using antidepressants was comparable to healthy controls, but the level of evidence was not conclusive (p(m_0_|Y) = 0.53). The analyses for effects on individual nodes were in line with the above (Figure 2). These findings were mostly robust against adjustments for potential confounding variables (i.e., age, sex, depression severity, presence of recurrent depression, presence of psychotic depression, use of lithium carbonate, use of antipsychotics, and site variables; SI Figure 4). The only substantial difference in the adjusted analyses appeared in patients using benzodiazepines, showing increased self-inhibition exclusively in specific network nodes rather than a widespread increase across networks.

**Figure 2.**
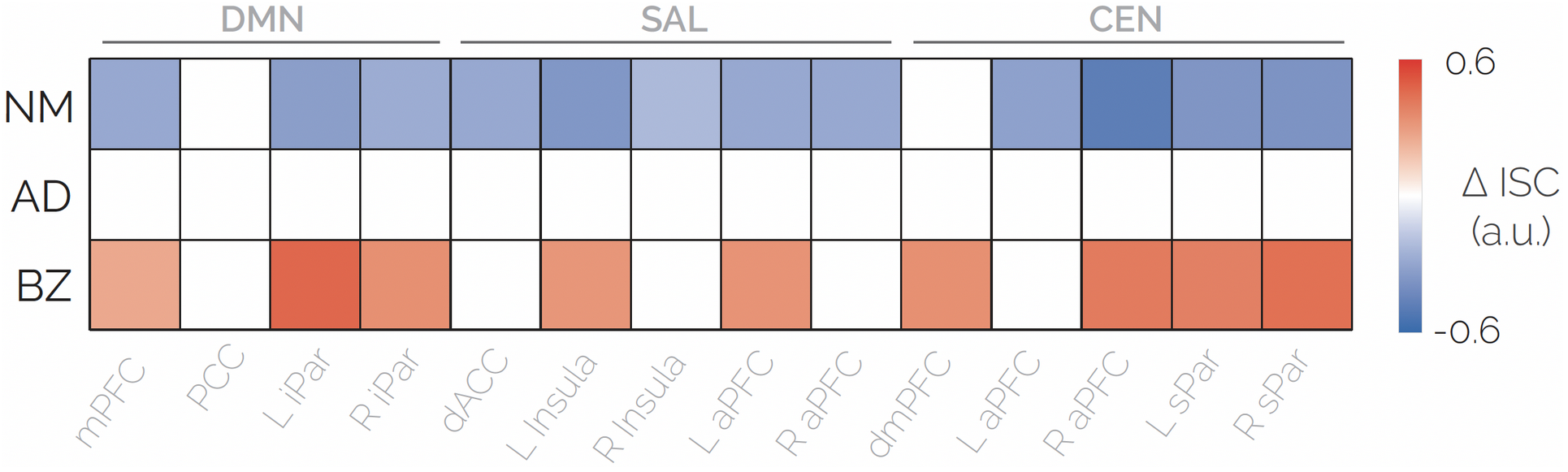
Group differences in region-specific inhibitory self-connections. Results of the analysis using automatic search over reduced models to obtain probabilities for specific connectivity parameters. Colored squares indicate very strong evidence (*p*(Y|m) > 0.99) for increased (red) or reduced (blue) self-inhibition between the specified patient group and healthy controls. ΔISC = difference in inhibitory self-connection strength; AD = patients using antidepressants; BZ = patients using benzodiazepines NM=patients not using AD or BZ. DMN = default mode network; SAL = salience network; CEN = central executive network. L= left; R = right; mPFC = medial prefrontal cortex; PCC = posterior cingulate cortex; iPar = inferior parietal cortex; dACC = dorsal anterior cingulate cortex; aPFC = anterior prefrontal cortex; dmPFC = dorsomedial prefrontal cortex; sPar = superior parietal cortex.

We used post-hoc pairwise comparisons to test for differences between patient groups (SI Methods and SI Figure 5). Compared to unmedicated patients, patients using antidepressants and/or benzodiazepines showed increased inhibitory self-connection strengths in all nodes across networks (p(m_4_|Y) = 1.0 for both groups). Compared to patients using antidepressants, benzodiazepines users showed increased inhibitory self-connections in specific nodes (particularly in the CEN; SI Figure 5). The analyses were adjusted for the same covariates as in the group comparison model described above and adjusted for the number of post-hoc comparisons.

In sum, these results yielded that, compared to unmedicated patients, antidepressants users showed a higher degree of self-inhibition which was more similar to healthy controls. Importantly, even though inhibitory deviations (from healthy controls) seemed to be normalized in patients using antidepressants, all patient groups still suffered from severe depressive symptoms (Table 1).

Next, an analysis was conducted on associations between inhibitory self-connections and depression severity in the distinct patient groups, as measured by the Hamilton Depression Rating Scale (HAM-D^29^). The results showed that more severe depression was associated with reduced self-inhibition across networks in unmedicated patients (p(m_4_|Y) = 1.0; SI Figure 6). Except from a few individual nodes, no clear associations between depression severity and inhibitory self-connections were found in patients using antidepressants and/or benzodiazepines (p(m_0_|Y) = 0.51 and p(m_0_|Y) = 0.43, respectively). The results were robust against adjustments for potential confounders (i.e., age, sex, presence of recurrent depression, presence of psychotic depression, use of lithium carbonate, use of antipsychotics, and site variables; SI Figure 6).

### Pharmacotherapy-related effects of excitation/inhibition imbalance on ECT effectiveness

Next, we examined whether excitation/inhibition imbalance at baseline was associated with ECT effectiveness. We first extracted the inhibitory self-connection parameters of all ROIs and identified the first principal component as a summary measure for regional self-inhibition of cortical networks throughout the brain. Using multiple regression models, we associated this summary measure to ECT effectiveness (i.e., the change in HAM-D scores from before to after ECT). The analysis was adjusted for baseline HAM-D scores and other known predictors of ECT effectiveness (i.e., presence of psychotic features, age, number of ECT-sessions with bilateral electrode positioning, and site variables; see SI Methods). The results showed a quadratic association between inhibitory self-connection and HAM-D change in the total sample (F(1,143) = 16.2, *p* < 0.001, Cohen’s *f*^2^ = 0.11; Figure 3a). Further analyses in the subgroups revealed that the quadratic relation was explained by medication related differences. Whereas no linear association was found in currently unmedicated patients (F(1,52) = 0.72, *p* = 0.40), patients using concomitant benzodiazepines and/or antidepressants showed a negative linear association between inhibitory strength and HAM-D change (F(1,47) = 11.58, *p* = 0.001 for antidepressants users; F(1,56) = 11.50, *p* = 0.001 for benzodiazepines users; see Figures 3b and 3c). The associations between neural inhibitory strength and ECT effectiveness in patients using pharmacotherapy showed a medium effect size (Cohen’s *f*^2^ = 0.21 for benzodiazepines users; Cohen’s *f*^2^ = 0.25 for antidepressants users), indicating that excessive inhibitory strength might be an important predictor of ECT non-response. Since the regression models were adjusted for potential confounders, the degree of variation in ECT effectiveness explained by inhibition seemed to be independent from other known predictors.

**Figure 3.**
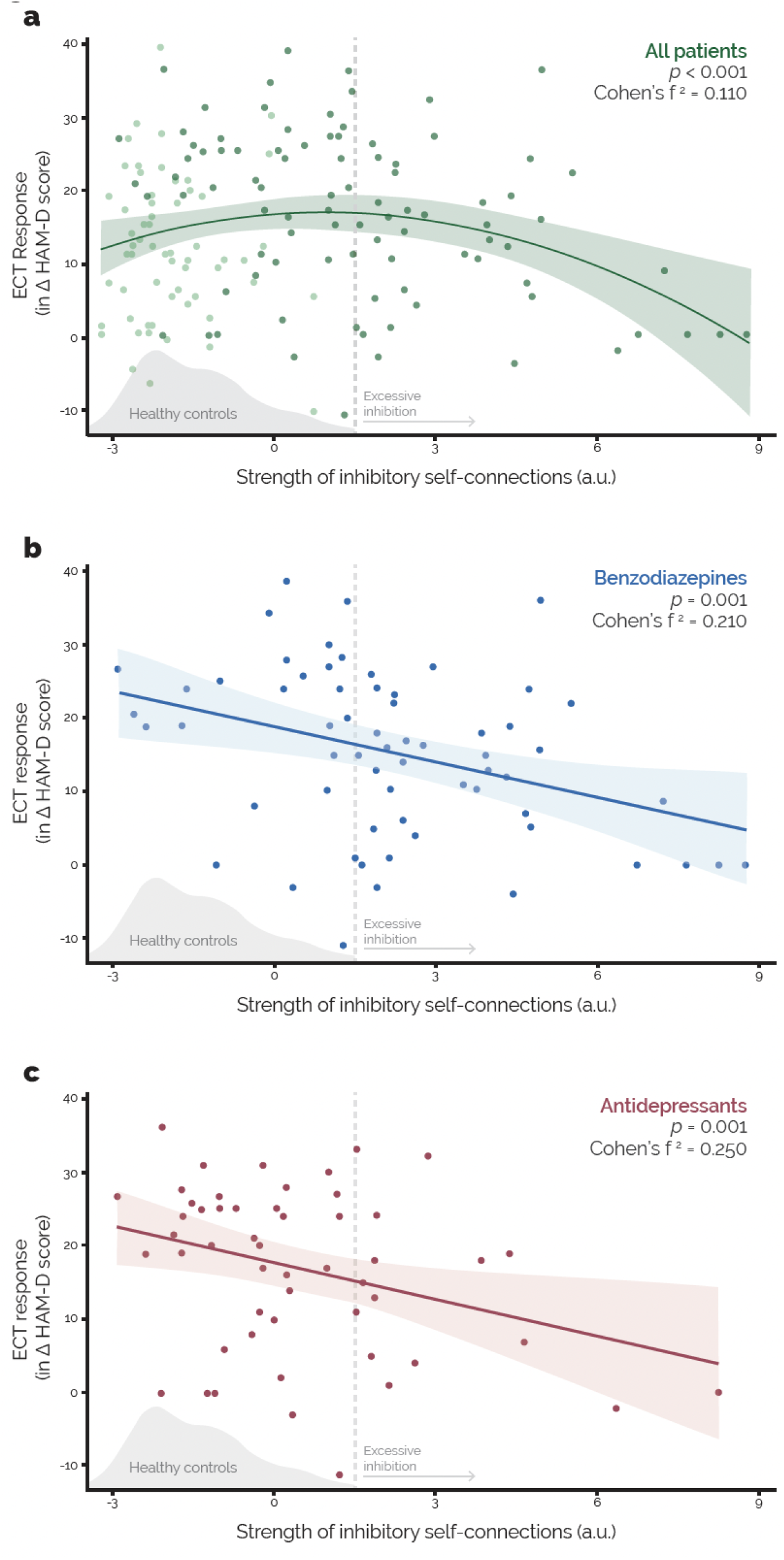
Associations between baseline inhibitory self-connection strength and ECT effectiveness. Regression plots on the association between the first principal component of inhibitory self-connections (arbitrary units) and the antidepressant effectiveness of ECT (change in HAM-D score before and after the ECT-course). The figure shows the regression plot for all patients (a), including both unmedicated patients (lighter green) and medicated patients (darker green). Additionally, separate regression plots are shown for patients using benzodiazepines (b) and patients using antidepressant (c). For each regression analysis, p-values and effect sizes for the displayed association are included adjusted for covariates. A medium effect size is indicated by 0.15 ≤ *f*^2^ ≥ 0.35. In each plot, a density plot is included to show the distribution of self-inhibition in healthy controls. The strength of patient’s self-inhibition that appeared beyond the range of healthy controls is identified as excessive.

For the analysis of clinical data, we used a multiple regression model to examine the association between relevant predictors — including antidepressants and benzodiazepines use — and ECT effectiveness (see SI Methods). In line with previous research^30^, the analyses revealed that the use of antidepressants enhanced ECT effectiveness (F(1,158) = 7.32, *p* = 0.008, Cohen’s *f*^2^ = 0.05). Therefore, our findings regarding the negative effects of inhibitory strength on ECT effectiveness mediated by antidepressants use seemed counterintuitive. Further analyses showed that the use of medication enhanced ECT effectiveness as long as the inhibitory strength was within the range of healthy controls (Figure 3b and 3c). This appeared true for both patients using antidepressants (mean ΔHAMD = 18.2±11.1 SD; t(75) = 3.02, *p* = 0.003) and benzodiazepines (mean ΔHAMD = 19.6±12.5 SD; t(37) = 2.83, *p* = 0.007), compared to unmedicated patients (mean ΔHAMD = 11.6±10.0 SD). However, excessive inhibition (defined as an inhibitory strength beyond the range of healthy controls) was present in 27.8% of antidepressants users and 60.3% of benzodiazepines users. None of the unmedicated patients showed excessive inhibition. Regarding the change in HAMD-scores, patients that showed excessive inhibition while using antidepressants (mean ΔHAMD = 12.5±9.7 SD; t(22) = 0.30, *p* = 0.765) and/or benzodiazepines (mean ΔHAMD = 12.3±9.6 SD; t(81) = 0.33, *p* = 0.738) did not differ from patients without concomitant medication use. This indicated that the positive effects of concomitant pharmacotherapy on ECT effectiveness were nullified if the degree of self-inhibition was excessive.

### Influence of ECT on excitation/inhibition balance and associations with clinical effectiveness

We used a longitudinal design to examine changes in excitation/inhibition ratios after the ECT-course, compared to baseline, and its associations with antidepressive effectiveness. We used a multisession resting-state spDCM analysis in a subsample of participants for whom longitudinal data were available (n_total_ = 113; n_patient_ = 89; n_HC_ = 24). Patients were scanned before and after the ECT-course. Healthy controls were also scanned twice, approximately four weeks apart. First, a model was run to test for effects of ECT on inhibitory self-connections in patients compared to longitudinal effects in controls. This model also tested for associations between change in inhibitory self-connections and treatment effectiveness (i.e., change in HAM-D; SI Figure 7). The analysis yielded weak evidence suggesting that longitudinal effects of ECT in patients did not differ from longitudinal effects in healthy controls (p(m_0_|Y) = 0.67). The model did yield conclusive evidence showing that ECT effectiveness was not associated with changes in inhibitory self-connections in any of the networks (p(m_0_|Y) = 0.99). These results were robust against adjustments for potential confounding variables (i.e., age, sex, baseline depression severity, presence of recurrent depression, presence of psychotic depression, lithium carbonate use, antipsychotic use, number of ECT-sessions, and site variables). Also, analysis on ECT response (i.e., a binary variable indicating >50% reduction of HAM-D scores after ECT) yielded very similar results (SI Figure 8). Because the patient groups (defined by medication use) showed different baseline inhibitory self-connection strengths, we tested whether longitudinal effects differed between groups. The results showed that ECT effectiveness was not associated with changes in inhibitory self-connections in any of the patient groups (p(m_0_|Y) = 1.0 for each group; SI Figure 9).

## Discussion

In this study, we investigated the role of the excitation/inhibition imbalance in the treatment of severe depression. The cross-sectional results revealed that unmedicated patients showed reduced inhibition compared to healthy controls. Also, more severe depression was associated with reduced self-inhibition in this group. Patients using antidepressants and/or benzodiazepines showed increased inhibition compared to unmedicated patients. The degree of inhibition in patients using antidepressants seemed comparable to healthy controls, whereas patients using benzodiazepines showed increased inhibition compared to controls. In these medicated patients, the degree of inhibition was not associated with depression severity. In line with previous findings^12–14^, these findings suggest that pharmacotherapy reverses the reduced inhibition that is associated with depression.

However, patients in this study all suffered from perpetuating severe depressive symptoms, indicating that the reversal of the excitation/inhibition imbalance in patients using pharmacotherapy, in and of itself, did not result in a persistent antidepressive effect. Additionally, the subsequent ECT effectiveness was not associated with changes in the strength of self-inhibition. Finally, we report that many patients using pharmacotherapy showed a degree of inhibition beyond the range of healthy controls. If the degree of inhibition was within the healthy range, concomitant antidepressants and/or benzodiazepines use was associated with increased ECT effectiveness. Remarkably, the presence of pre-treatment excessive neural inhibition nullified these positive effects of concomitant pharmacotherapy on the outcome of ECT.

An influential hypothesis poses that deficits in neural excitation/inhibition balance represent a fundamental aspect of depression and that reversing these deficits may be an important property to the antidepressive effect of existing treatments^6–9^. In line with this hypothesis, unmedicated severely depressed patients showed reduced inhibitory selfconnections compared to healthy controls. But contrary to this hypothesis, our results suggest that reversal of inhibitory deficits by using antidepressants and/or benzodiazepines is, in and of itself, not a sufficient condition for an antidepressive treatment effect in patients with severe depression. This indicates that either the effectiveness of pharmacotherapy in severe depression did not depend on reversing the inhibitory deficits, or that additional conditions were needed for an antidepressant response (i.e., the absence/presence of other biological, psychological, or social phenotypes). One of such additional conditions could be that an increase in inhibition should be accompanied by an increase in excitation for an antidepressant effect^31^. Also, changes in self-inhibition were not associated with the effectiveness of subsequent ECT, indicating that reversal of the excitation/inhibition balance neither was a necessary condition for the effective treatment of severe depression. Based on these findings, the previously stated hypothesis might be partially altered. Although the excitation/inhibition imbalance still seemed a key aspect of the neural etiology of severe depression, its reversal by treatment might not be a sufficient nor a necessary condition for an antidepressive effect.

Recent studies showed antidepressant effects of a positive allosteric modulator of GABA_A_ receptors in post-partum depression^32^ and major depressive disorder^33^. In contrast to our findings, these findings suggest that reversing the inhibitory deficit would be a sufficient condition for an antidepressant effect in severe depression. However, the results of these trials may reflect antidepressant effects that are not maintained at the longer-term. The antidepressive effectiveness in the trial on major depressive disorder vanished within six weeks after initiating the pharmacotherapy^34^, and the trial on post-partum depression used a follow-up period of only four weeks. Such short-term antidepressant effects are also observed in the treatment of depressed patients with benzodiazepines, a well-known GABA_A_ modulator^15^. In our study, data on the duration of current pharmacotherapy use was not available. However, severely depressed patients indicated for ECT are typically characterized by long-term pharmacotherapy use (i.e., at least multiple months, but most likely years). Therefore, our findings probably reflect the clinical and neural state of patients in a later phase of pharmacotherapy, as opposed to the early phase studied in the mentioned studies^32,33^. We therefore hypothesize that, even though reversing the excitation/inhibition imbalance might induce short-term antidepressant effects, it might not be sufficient for a long-term clinical response.

Our findings on the effects of pharmacotherapy on the excitation/inhibition balance might be specific to more severe and treatment-resistant subtypes of depression and its treatments. Preliminary evidence suggests that severe depression seems to differ from milder subtypes of depression with respect to GABA-ergic dynamics^35,36^. For instance, a transcranial magnetic stimulation (TMS) study indicated that GABA_B_ receptors seem to be affected by all subtypes of depression, whereas changes in GABA_A_ receptors might be unique to treatment-resistant depression^37^. Therefore, the pharmacotherapy-related effects on the excitation/inhibition imbalance in our study might be established by changes in specific GABA receptor types or subunits that were insufficient for achieving effectiveness in these treatment-resistant patients, while changes in other receptor types or subunits might. However, the lack of an association between ECT effectiveness and changes in self-inhibition suggests that changes in GABA-ergic dynamics are not necessary for an antidepressant response in severe depression. Also, in other subtypes of depression, reversing the excitation/inhibition imbalance may be sufficient for an antidepressive response. Therefore, further research is needed to test whether our findings generalize across other subtypes of depression.

We also provide evidence suggesting that an optimized excitation/inhibition balance at baseline is associated with increased ECT effectiveness, whereas too little or too much inhibition can result in a lower antidepressive response. One possible explanation of this effect is that excessive inhibition due to pharmacotherapy may reduce the propagation of the induced seizure. For decades, it has been suggested that seizure propagation between distant regions of the brain is pivotal for ECT effectiveness, including cortical–thalamocortical interactions and direct cortical–cortical connections^19^. Consequently, excessive inhibition might obstruct spreading of seizure activity to essential parts of the patients’ brain. In support of this, a study using TMS showed that baclofen, a GABA_B_ receptor agonist, reduced interhemispheric signal propagation^38^. Also, previous studies showed that the strength of the post-ictal suppression is associated with ECT effectiveness^21,39^. Post-ictal suppression is an electroencephalographic phenomenon characterized by an ictal slow-wave phase, associated with inhibitory restraints, after which seizure activity is terminated. The interhemispheric coherence in ictal slow-wave activity has been associated with greater ECT effectiveness^21,40^, indicating that individual differences in the seizure termination processes may explain the variation in treatment effectiveness between patients. These termination processes may be suboptimal in patients if the degree of self-inhibition is reduced. In light of the above, our findings denote that an optimal excitation/inhibition balance at baseline increases the effectiveness of ECT by allowing for a proper propagation and termination of the induced seizure.

ECT did not seem to affect the strength of neural inhibition in our patients, when compared to changes in inhibition over time in healthy controls (although evidence was weak). Some magnetic resonance spectroscopy (MRS) studies showed increase of inhibitory GABA concentrations over the ECT-course, but adding healthy controls diminished these significant findings^41^. Additionally, we provide strong evidence that ECT effectiveness did not depend on changes in regional self-inhibition. This is compatible with previous null findings in studies using frequentist statistics^41^. However, the classical statistics used in these studies were not able to provide conclusive evidence on the absence of an association between GABA concentrations and ECT effectiveness. Using a Bayesian approach, we provide very strong evidence indicating that the effectiveness of ECT did not depend on changes in the excitation/inhibition balance^42^.

In our study, use of pharmacotherapy influenced the relation between neural inhibition and ECT effectiveness. In daily practice, combining ECT with antidepressants is regularly attempted to further enhance effectiveness^43^. Also, a recent meta-analysis of randomized controlled trials showed that the use of antidepressants (i.e., as a dichotomous variable) increased ECT effectiveness when considering the entire group of patients^30^. However, based on our findings, this statement may be nuanced by adding that the positive effects of antidepressants can be nullified if the degree of inhibition (associated with pharmacotherapy use) is excessive. The strength of neural inhibition in medicated patients is probably due to a combination of pharmacotherapy dosage effects and variability in drug metabolism between patients^44^. Consequently, the administered dosage of concomitant antidepressants and, in particular, benzodiazepines may affect the outcome of ECT. This may also explain the contradictory results in the current literature, in which studies reported positive, negative and absence of effects of concomitant benzodiazepines use on ECT response^45^. Additionally, higher dosages of concomitant used benzodiazepines (i.e., lorazepam) were associated with shorter seizure durations during ECT^46^. To mitigate these effects, the competitive benzodiazepine-antagonist flumazenil can be used. Although, it was not known whether or which patients received flumazenil in our study, this could not prevent that patients with excessive inhibition beforehand showed lower ECT effectiveness. Thus, based on our findings we hypothesize that, while the use of concomitant pharmacotherapy (as a dichotomous variable) may enhance ECT effectiveness, higher dosages may result in a poorer clinical response. But, to provide conclusive evidence on this hypothesis, randomized clinical trials have to be conducted, to test dose-dependent (and preferably plasma-level controlled) effects of benzodiazepines and antidepressants on ECT effectiveness. To the best of our knowledge, no such trials have been reported in the literature yet.

Our findings may guide future research on treatments for severe depression. The findings suggested that reversing the excitation/inhibition imbalance by pharmacotherapy might not be sufficient in severe depression. This may inform the direction of drug development for depression, especially regarding agents targeting GABA. Additionally, we identified excessive inhibition as a potential novel biomarker of ECT ineffectiveness in patients using benzodiazepines or antidepressants. In our study, the effect size of this biomarker was comparable to the presence of psychotic features (data not shown), which is currently one of the most powerful predictors of ECT outcome^47^. Another advantage of this biomarker is that it may be modifiable by adjusting the pharmacotherapy dosages in the individual ECTpatient. Thereby, our findings suggest that pharmacotherapy management informed by the excitation/inhibition ratio may prevent excessive inhibition and potentially enhance the effectiveness of ECT. Alternatively, patients treated with ECT may simply benefit from lower dosages of pharmacotherapy during the ECT-course. This may especially hold for patients using benzodiazepines, which showed higher risks of excessive inhibition in this study. However, replication neuroimaging and clinical studies are needed before implementation of these findings in clinical guidelines.

Several limitations have to be taken into account when interpreting our findings. First, the effects of pharmacotherapy on the excitation/inhibition balance were studied using cross-sectional data and an observational design. These findings should be replicated in longitudinal prospective studies to rule out potential selection bias and effects of confounding variables not tested in the current study (e.g., comorbid psychiatric disorders, level of anxiety, concomitant use of other medication). Also, data on the timing of pharmacotherapy (dis)continuation was unavailable, which should be taken into account by future studies. Additionally, effects of pretreatment with flumazenil could not be evaluated. Furthermore, our study was observational and included severely depressed patients that failed to respond to pharmacotherapy and were indicated for ECT. This is a highly selected and severely ill population to test the general neural mechanisms of pharmacotherapy. Therefore, our findings may not necessarily generalize to other subtypes of depression. Moreover, we established neural inhibition/excitation on the level of cortical networks, each node containing millions of neurons. This restrained our level of inference and made conclusions on specific cellular processes impossible (e.g., neurotransmitter concentrations, effects on specific receptor types or subunits). Finally, no (sufficient) data was available to test other effects of interest, such as dose-dependency of pharmacotherapy effects. Future studies have to be conducted to overcome these limitations.

In conclusion, our study provides novel evidence on the role of excitation/inhibition imbalance in treatments for severe depression. We show that reversing the excitation/inhibition imbalance may not be sufficient nor necessary for treatment effectiveness using pharmacotherapy or ECT in severe depression. Furthermore, the presence of excessive neural inhibition nullified the positive effects of concomitant pharmacotherapy on the antidepressant effectiveness of ECT. These findings should be replicated by future studies to assess generalizability to other samples and milder subtypes of depression. Nonetheless, this work provides a modest but important step for understanding the working mechanisms of treatments for severe depression and enhancing their effectiveness.

## Methods

### Participants and treatment

Data were obtained from the Global ECT-MRI Research Collaboration (GEMRIC)^27^. All patients in this study were indicated for ECT and were treated according to internationally accepted guidelines. The data used for this study were acquired at seven sites across Europe and North America. Participants were divided into four groups: healthy controls (n = 59), treatment-resistant patients without current use of antidepressants and benzodiazepines (noAD/BZ; n = 60), treatment-resistant patients using antidepressants (AD; n = 56), and treatment-resistant patients using benzodiazepines (BZ; n = 63). The latter two groups showed considerable overlap (n = 24), which was accounted for in the analyses. All patients had a clinical diagnosis of unipolar major depressive disorder, classified according to the International Statistical Classification of Diseases and Related Health Problems (ICD-10). Patients without current use of benzodiazepines and antidepressants did receive at least one trial of pharmacotherapy before. Descriptive data on demographics (age, sex) and clinical variables (depression severity, presence of psychotic features, presence of recurrence) were available. Depression severity was assessed using the Hamilton Depression Rating Scale (HAM-D) or Montgomery-Åsberg Depression Rating Scale (MADRS), depending on the sites’ preferences. Ratings of the MADRS were converted to HAM-D scores, using the following formula^48^:

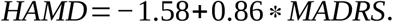

Patients were excluded if (1) bipolar disorder was present^49^, (2) BZ or AD use data were incomplete, and (3) if neuroimaging data were unavailable or of insufficient quality.

The ADs that were examined in this study consisted of selective-serotonin reuptake inhibitors (SSRIs), selective-noradrenaline reuptake inhibitors (SNRIs), tricyclic antidepressants (TCAs) and monoamine oxidase inhibitors (MAOIs). In the group of patients currently using AD, 34% used SSRIs, 36% used SNRIs, 27% used TCAs and 3% used MAOIs. The specific type of BZ, and dosages for AD and BZ, were not available in the database. Regarding used electrode placements, 126 patients received ECT with right unilateral (RUL) and 62 patients with a bilateral (BL) electrode placement. The average number of ECT-sessions per course was 12.4 ± 5.5 SD.

### fMRI acquisition, preprocessing, and quality control

Structural and functional resting-state magnetic resonance imaging (MRI) data were acquired at seven sites (for scanning parameters per site, see SI Table 4). The fMRI preprocessing steps were identical for each participant. First, structural and functional images were reoriented and brain extraction was performed using Advanced Normalization Tools (ANTs v2.2.0). Images were coregistered using ANTs and FMRI Software Library (FSL v5.0.10) using boundary-based registration. After discarding the first two volumes for each fMRI series, FSL’s MCFLIRT was used to apply head movement correction by realigning the volumes to the middle volume using six parameters for rigid body transformations. Functional images were spatially smoothed using a Gaussian kernel with 5 mm full-width at half-maximum. For motion correction, an ICA-based strategy for Automatic Removal of Motion Artifacts (ICA-AROMA) was used. The noise components estimated by ICA-AROMA were used to compute denoised cosines for high pass filtering (f=0.009). Furthermore, mean white matter (WM) and cerebrospinal fluid (CSF) timeseries were computed as additional nuisance variables. Temporal high-pass filtering was used to remove low-frequency drifts (< 0.01 Hz) and images were registered to 4 mm isotropic voxel size. Both the denoised cosines and WM/CSF nuisance variables were used to denoise the fMRI data and perform high pass filtering in one single step^50^ ANTs were used for normalization of transformation matrices to MNI space using 2 mm standard templates.

After the preprocessing, a quality control procedure was performed. First, participants that showed a high degree of motion were identified using three criteria: (1) any rotation/translation exceeding 4 mm/degrees, (2) the average framewise displacement (FD) exceeding 0.3 mm, and (3) less than 4 minutes of RS-fMRI data unaffected by motion (FD<0.25mm). If any of the criteria applied to a participant’s RS-fMRI data, the participant was excluded from the study. Then, the data were visually inspected to confirm high-quality coregistration and normalization of the T1w and RS-fMRI scans. Finally, the EPI signal-to-noise and field of view were assessed to identify potential signal dropout or other artifacts. Again, participants with low quality preprocessed data were excluded from the analysis. Finally, RS-fMRI of sufficient quality was available for a total of 214 participants with respect to cross-sectional baseline data. Regarding longitudinal data, RS-fMRI data of sufficient quality was available for 113 participants.

### First-level spDCM analysis

Spectral dynamic causal modeling (spDCM) for RS-fMRI was used to estimate effective connectivity parameters^22^. Analyses were conducted in Statistical Parametric Mapping (SPM12, revision 7771) with DCM12.5 (revision 7497). spDCM used a generative forward model with two components. The first one described how neuronal populations causally interact. The second component mapped neuronal activity from the first model to observed (crossspectra of) hemodynamic responses. Thereby, spDCM was able to estimate directed connectivity between brain regions as well as regional self-connections. The construct validity of spDCM has been shown by its ability to accurately recover effective connectivity in simulated data^22,23^, and specific effective connections corresponding to well-documented neural pathways in rodents^24^. The self-connections were assumed to be inhibitory, which was in line with in vivo animal studies^25^. By regulating the strength of inhibition, the self-connections allowed for sustained activity in the cortex while maintaining a balance between excitation and inhibition throughout the network. A recent magnetoencephalography study using DCM showed that a GABA_A_ reuptake inhibitor altered the strength of inhibitory (self-)connections in humans^26^, indicating that the DCM framework could be used to study the effect of therapeutic interventions on (self-)inhibition.

We used the following procedure. First, the RS-fMRI timeseries were extracted from fourteen regions of interest (ROIs). These regions formed the core of three networks (i.e., default mode network (DMN), salience network (SAL) and central executive network (CEN)) that are of central importance to depression^28^ and psychopathology in general^51^. Time series were extracted from a sphere with 5 mm radius centered around MNI coordinates extracted from previous research (see SI Figure 1)^52^ single representative time series was established by retaining the first component of a principal components analysis across voxels in each sphere. For baseline analysis, a full spDCM was estimated for each participant individually (i.e., all 196 connections in matrix A of the neural model, henceforth the ‘A-matrix’, were turned on). Default settings were used for the individual-level spDCM analyses and no experimental inputs were specified (i.e., all 196 connections in the ‘B-matrix’ turned off). The expected values and covariances of the estimated parameters of the ‘A-matrix’ were then taken to the group-level analysis.

For the longitudinal analysis, a multisession resting-state spDCM approach was used. Only the longitudinal effect of ECT on inhibitory self-connections were estimated. Therefore, a full ‘A-matrix’ was specified and an identity matrix was used as the ‘B-matrix’. In this case, the ‘A-matrix’ provided the averaged connectivity over both sessions and the ‘B-matrix’ modelled the change between sessions. Because we were interested in changes associated with ECT, the estimated parameters of the ‘B-matrix’ were taken to the group-level analysis.

### Parametric Empirical Bayes

Group-level effects could be estimated using the Parametric Empirical Bayesian (PEB) framework^53^. A PEB general linear model (GLM) was able to infer effects at multiple levels (e.g., group effects or between-session effects) by combining the expected values and covariances from all individual-level spDCM parameters. Additionally, the PEB model was able to finesse spDCM parameters at the individual level by using group-level posteriors as empirical priors^54^. PEB models were used for studying associations between baseline self-inhibition and ECT effectiveness (see Analysis on baseline associations with ECT effectiveness, below).

For each PEB model (see below for PEB model designs), effects on inhibitory selfconnections in individual nodes and effects on combinations of self-connections of multiple nodes were examined. For effects on individual self-connections, Bayesian Model Reduction (BMR) was used to automatically search over reduced models with certain connectivity parameters ‘turned off’ (i.e., fixed at their prior mean of zero). In this method, all possible reduced models were compared, and models with connectivity parameters that did not contribute to model evidence were iteratively discarded. This process was continued until removing parameters reduced the model evidence. Subsequently, a Bayesian model average was calculated over the 256 models with the largest model evidence in the final iteration (or over the remaining models, if less than 256 models survived). Using this ‘pruning’ and averaging approach allowed inference on the posterior probability of specific connectivity parameters. A threshold for posterior probabilities indicating strong evidence was used for each second-level parameter (p(m|Y) > 0.95).

For the analysis on specific groups of connections, BMR was used to estimate the model evidence for pre-specified reduced connectivity models. Subsequently, the model evidence between the reduced models was compared to that of an empty model (i.e., all parameters ‘turned off’) to determine which reduced model best fitted the observed data^54^. Five pre-specified reduced models were used, in which models included self-connections in none of the networks, one of the networks (DMN, CEN or SAL), or all of the networks (DMN, CEN and SAL; see SI Figure 3). Thereby, this procedure tested whether associations between variables of interest (e.g., depression severity) and self-connections parameters were absent, network-specific or widespread throughout brain networks.

Again, a threshold of a posterior probability indicating (at least) strong evidence in favor of a model was used (*p*(m|Y) > 0.95).

### PEB model designs

Multiple PEB GLMs were run to test for group differences in inhibitory self-connections at baseline, associations with depression severity, and changes in inhibitory self-connections before and after ECT.

First, an analysis on group differences between healthy controls and patient groups was conducted. Group differences could be modeled using multiple GLM designs. We used a model selection procedure by comparing the two most convenient designs with respect to model evidence. A GLM in which all patient groups (unmedicated, AD, BZ) were modeled with a separate binary regressor (and healthy controls as the intercept, i.e., reference group) showed greatest model evidence, and was selected for the analysis on group differences (see SI Figure 2 for the models and difference in model evidence). Additionally, an identical model with extra regressors was run to test if the results of the initial model were robust against effects of potential confounding variables. The added regressors consisted of age, sex, baseline HAM-D score, presence of recurrent depression, presence of psychotic depression, lithium carbonate use, antipsychotic use and site variables. The age and sex variables were mean centered, such that the reference comparison group (i.e., the intercept) represented healthy control subjects of a group average age, averaged over males and females. Due to the collinearity constraint in a GLM design, not all sites variables could be included as covariates (i.e., the sum of all site variables equals the intercept). Therefore, site variables that added substantially to the model evidence were selected as potential confounders. These variables were selected by comparing the model evidence between the model with demographic and clinical covariates with an identical model plus one of the site variables. If a site variable explained considerable variation in the data, the model fit increased. However, adding a covariate also added extra complexity to the model, which reduced model evidence. This procedure tested whether the increase in the goodness-of-fit by adding the site variable outweighed the increase in complexity. Site variables that increased model evidence were included as a potential confounder in the GLM (SI Figure 3).

Because the model above did not allow direct comparisons between patient groups, post-hoc pairwise comparisons between the AD, BZ and unmedicated patient groups were conducted. For each comparison, a GLM was created that included patients of the specified patient groups (see SI Figure 5 for model design). The same potential confounding variables as in the model above were also included in the model. Additionally, AD use was included in the comparison between patients using BZ and unmedicated patients, and BZ use was included in the comparison between patients using AD and unmedicated patients. The latter was done to account for the overlap between the AD and BZ groups. For post-hoc analyses, a threshold that indicated strong evidence according to Bayesian statistics was used, adjusted for multiple comparisons using Bonferroni (p(m|Y) > 0.983).

Associations with depression severity were analyzed using a GLM that included patients only (see SI Figure 6 for the model design). This model tested if there were effects of depression severity on inhibitory self-connections in the complete group (i.e., main effect of baseline HAM-D) or specific patient groups (i.e., patient group by baseline HAM-D interactions). Aside from the unadjusted model, a model was run to test if the results were robust against potential confounders. The same demographic and clinical covariates were included as in the adjusted model for group comparison. Also, an identical procedure was used to select relevant site variables (i.e., selecting site variables that increased model evidence).

Finally, PEB models were used to test for longitudinal effects of ECT. Parameters of the first-level longitudinal analyses were entered into group-level PEB models. The first GLM tested for longitudinal effects in patients compared to healthy controls, and for associations with ECT effectiveness (SI Figure 7). Again, we subsequently tested if the results were robust against potential confounding variables. Based on previous studies^47,55^ and clinical practice, the following demographic and clinical variables were included in the GLM as potential confounders: age, sex, baseline HAM-D score, presence of recurrent depression, presence of psychotic depression, lithium carbonate use, antipsychotic use, number of ECT-sessions, and site variables. Site variables were selected using an identical procedure as described in the analysis on group differences. Furthermore, an additional model was run for effects of treatment response (i.e., a binary regressor indicating HAM-D reduction greater than 50% compared to baseline), instead of the continuous measure for ECT effectiveness used in the model above. Finally, because patient groups showed different baseline strengths in inhibitory self-connections, we also tested for longitudinal effects of ECT in the distinct patient groups (SI Figure 9 for the model design). The latter model also tested if ECT effectiveness in distinct patient groups was associated with changes in inhibitory self-connections.

### Analysis on baseline associations with ECT effectiveness

For the clinical analysis, a multiple regression model was run to test the effect of clinical variables on ECT effectiveness, which was defined as the change in HAM-D scores before and after the ECT-course (HAM-D_PRE_ – HAM-D_POST_; i.e., the dependent variable). The included clinical variables were the use of BZ, the use of AD, baseline HAM-D scores, presence of psychotic features, age, number of ECT-sessions with BL electrode placement, and seven binary variables indicating site membership. These variables were selected because those were identified as predictors of ECT effectiveness in previous research^47^ and in clinical practice.

In order to relate the excitation/inhibition ratio at baseline to ECT effectiveness, multiple regression models were used in R^56^. A different approach was used for this analysis compared to the analyses described above (see PEB design models), because we aimed to provide effect size measures that were comparable to the existing literature on ECT outcome predictors (i.e., provided by classic frequentist statistics, rather than Bayesian statistics). First, a PEB GLM (with only the intercept) was used to establish the posterior probability distribution of each of the 14 self-connection parameters for each individual. The mean of the probability distribution for each self-connection was extracted for each participant, and the first principal component (as obtained by principal component analysis using the *prcomp* package) was used as summary measure of the excitation/inhibition balance. Subsequently, a classical (non-Bayesian) multiple regression model was used to examine the relation between excitation/inhibition balance (i.e., the independent variable) and ECT effectiveness. Both linear and quadratic effects of the excitation/inhibition balance on ECT effectiveness were tested. All variables that showed a significant effect in the above-described clinical analysis were included as a covariate in the model (i.e., baseline HAM-D score, age, presence of psychotic features, number of ECT-sessions with BL electrode placement, and two binary variables indicating site membership). For post-hoc analyses, identical models were built that only included patients of the separate groups (i.e., noAD/BZ, AD and BZ). The same covariates were entered into the model. Because the AD and BZ groups showed considerable overlap, BZ use was added as a covariate in model of the AD group, and AD use was added in the model of the BZ group. This was done to ensure that the effects in de AD and BZ group were independent from BZ and AD use, respectively. Cohen’s *f^2^* was used as a measure of effect size, in which 0.15 ≤*f*^2^ ≥ 0.35 indicates a medium effect size^57^.

## Acknowledgements

The authors are very grateful to all patients who participated in this study. Additionally, we would like to thank all professionals that contributed to treatment delivery and acquisition of the data.

## Author contributions

F.t.D., W.B., C.A., M.A., A.D., L.E., P.F.P.v.E., E.v.E., P.C.R.M., K.N., I.T., D.R., P.S., M.V., J.V., M.v.V., H.B., L.O., J.A.v.W., & G.A.v.W contributed to data acquisition. F.t.D. primarily performed data analysis. W.B., P.Z., J.A.v.W., & G.A.v.W also made substantial contributions to data analysis. F.t.D., J.A.v.W. & G.A.v.W drafted the manuscript. W.B., P.Z, C.A., M.A., A.D., L.E., P.F.P.v.E., E.v.E., P.C.R.M., K.N., I.T., D.R., P.S., M.V., J.V., M.v.V., H.B., & L.O. substantially revised the manuscript.

