## Supplementary material for "Neural excitation/inhibition imbalance and the treatment of severe depression": Supplemantary figures

### Supplementary information

| Site | Scanner | Imaging type | Scanning parameters |
|---|---|---|---|
| Site 1 | Siemens Allegra 3T | T1 | Multi-echo MPRAGE, TR=2.53s, TE=1.74,2.6,5.46,7.32ms, Flip Angle=70, FoV=256x256 mm, voxel size=1.3x1.0x1.0mm |
| | | RS-fMRI | EPI TR = 2.0s, TE = 30ms, flip angle=70°, voxel size=3.4 × 3.4 × 5mm, 180 volumes |
| Site 2 | Philips Achieva 1.5T | T1 | Turbo field echo MRI, TR 7.6ms, TE=3.5ms, Flip angle=15, voxel size= 1.1x1.1x1.1mm |
| | | RS-fMRI | 2D gradient-echo single-shot EPI,TR 1.868s, TE=30ms, FoV=230mm, voxel size=2.4x2.4x4.5mm, 150 volumes |
| Site 3 | Siemens Avanto 1.5T | T1 | 3D MPRAGE, TR=2.25s, TE=3.68ms, flip angle=15, FoV=256x256x176, voxel size=1x1x1mm |
| | | RS-fMRI | EPI, TR=1.870s, TE=35ms, flip angle=80, FoV 224×224×137, voxel size = 3.5×3.5×3.0mm, 266 volumes |
| Site 4 | Siemens Trio 3T | T1 | Multi-echo MPRAGE, TR=2.53s, TE=1.64,3.5,5.32,7.22,9.08ms, flip angle=7, FoV=256x256x256mm, voxel size=1x1x1mm |
| | | RS-fMRI | Gradient-echo EPI, TR=2s, TE=29ms, flip angle=75, FoV=240x64x64mm, voxel-size=3.75x3.75x4.55mm, 154 volumes |
| Site 5 | GE Signa Hdxt 3T | T1 | 3D SPGR, TR=7.5 ms, TE=3 ms, FoV=240mm, voxel size=0.94x0.94x1mm |
| | | RS-fMRI | Gradient-echo EPI, TR=2s, TE=30 ms, FoV=240mm, Voxel size=3.75x3.75x3mm, 150 volumes |
| Site 6 | GE Signa Hdxt 3T | T1 | 3D SPGR, TR=7.84ms, TE=3.02ms, flip angle=12, FoV=256x256x256, voxel size=0.94x0.94x1mm |
| | | RS-fMRI | EPI, TR=1.8s, TE=35ms, flip angle=80, FoV=211mm, voxel size = 3.3×3.3×3.0mm, 202 volumes |
| Site 7 | Philips Intera 3T | T1 | TR = 9.6ms, TE = 4.6ms, flip angle=8, voxel size=0.98x0.98x1.2mm |
| | | RS-fMRI | EPI TR = 1.7s, TE = 33ms, flip angle=90, FoV= 230x128x230mm, voxel size=4x4x4mm, 250 volumes |

**SI Table 1 MRI acquisition parameters**

The parameters T1-weighted MRI and resting-state fMRI acquisition are presented for each site. TR = repetition time; TE = echo time; FoV = field of view; EPI = echo planar imaging; MPRAGE = magnetization prepared rapid gradient echo; SPGR = spoiled gradient recalled.

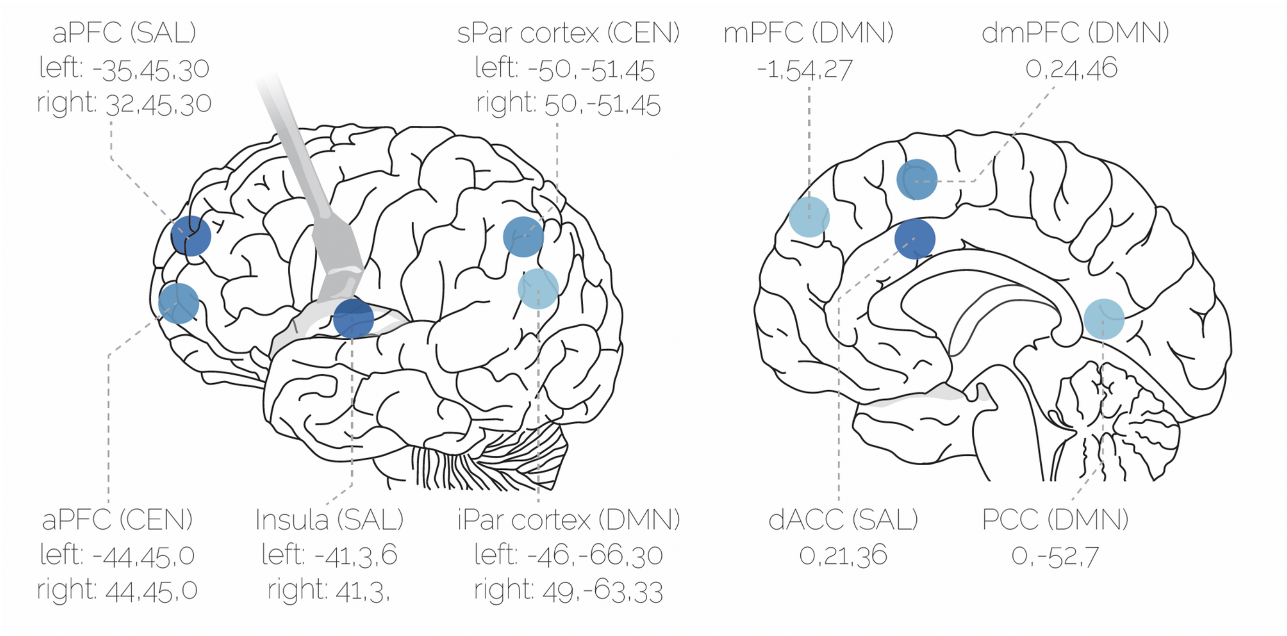


**SI Figure 1 Regions of interest**

Overview of the regions of interest that were included in this study with their respective MNI coordinates. The regions belong to three networks: the central executive network (CEN), salience network (SAL), and default mode network (DMN). aPFC = anterior prefrontal cortex; sPar = superior parietal cortex; mPFC = medial prefrontal cortex; dmPFC = dorsomedial prefrontal cortex; iPar = inferior parietal cortex; dACC = dorsal anterior cingulate cortex; PCC = posterior cingulate cortex.

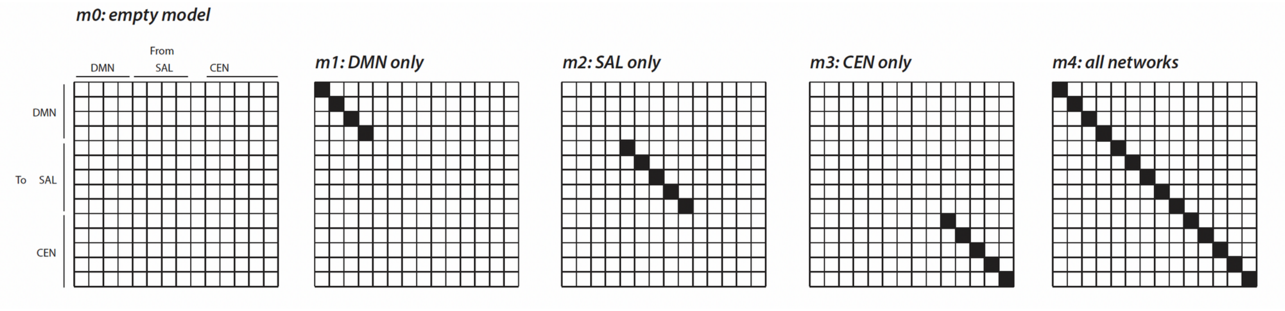


**SI Figure 2 Inhibitory self-connection models**

A schematic representation of the specific reduced models of inhibitory self-connections that were tested in the parametric empirical Bayes analysis. Models consist of a null-model (m0; testing for the evidence of absence), models for effects in specific networks (m1, m2, m3), a model for effects across networks (m4). Gray squares indicate that the connectivity parameter is ‘turned on’ and white squares indicate that the parameter is ‘turned off’ (i.e., fixed at their prior mean of zero). The model evidence was compared between models to determine which of these was the winning model for a specific parametric empirical Bayes analysis. Results are reported in the main text and SI Figures 4-9. DMN = default mode network; SAL = salience network; CEN = central executive network.

SI Fig 3

**Model 1**

$$\theta^{(1)} = HC * \theta^{(2)} + Pt * \theta^{(2)} + AD * \theta^{(2)} + BZ * \theta^{(2)} + \varepsilon^{(2)}$$

**Model 2**

$$\theta^{(1)} = HC * \theta^{(2)} + NM * \theta^{(2)} + AD * \theta^{(2)} + BZ * \theta^{(2)} + \varepsilon^{(2)}$$

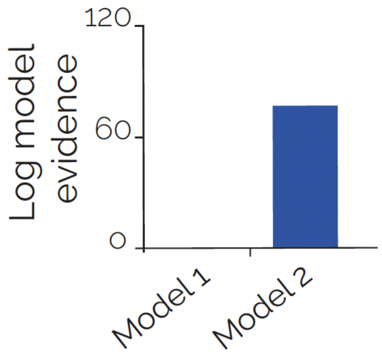


**SI Figure 3 Model selection for analysis on group differences**

To select the model design that best fitted the data, we compared two general linear models (GLMs) that tested for group differences regarding their log model evidence. The models described first level inhibitory self-connection parameters ($\theta^{(1)}$) by a linear combination of regressors and an error term ($\varepsilon^{(2)}$). Thereby obtaining group-level inhibitory self-connection parameters associated with each regressor ($\theta^{(2)}$). The change in log model evidence of model 2, relative to model 1, indicates that model 2 better fitted the data (right panel). HC = Healthy control; PT = all patients; AD = patient using antidepressants; BZ = patients using benzodiazepines; NM = currently unmedicated patients.

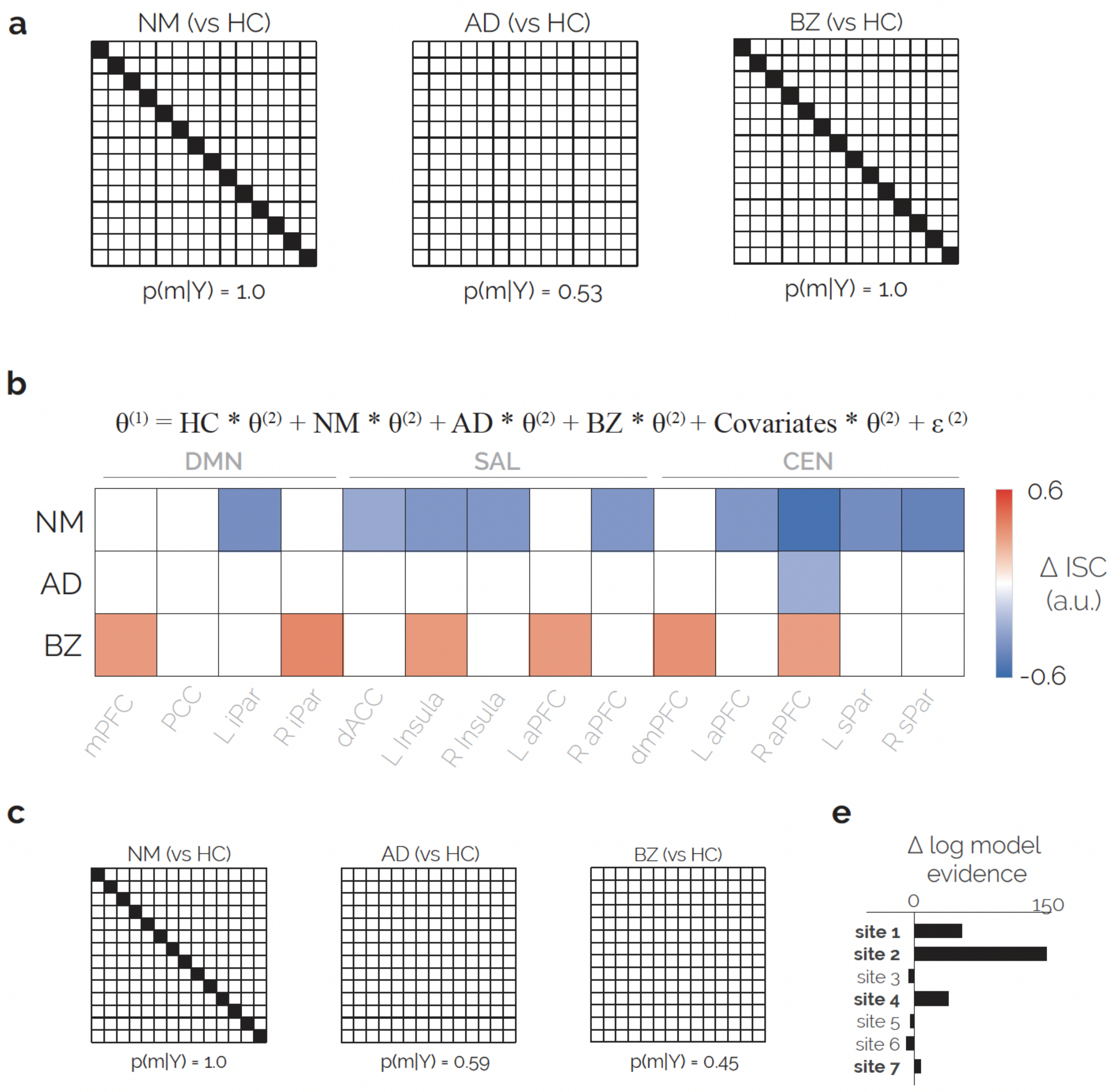


**SI Figure 4 Analyses on group differences**

Results on additional analyses on group difference of inhibitory-self connections. (a) For each patient group, compared to controls, the winning model is presented with its posterior probability. The results indicate that inhibitory self-connections across networks in the unmedicated and benzodiazepine group differed from healthy controls. Additionally, weak evidence for similarity between the antidepressant group and healthy controls is provided. Results on the automatic search over reduced models (i.e., for analysis on individual self-connections) are presented in Figure 2. (b) An additional analysis was performed to test whether results on group differences were robust against potential confounders. The model design (identical to model 2 in SI Figure 3 but with covariates) and results on the covariate-adjusted automatic search over reduced models are presented. (c) Results of the adjusted analysis on pre-specified models is presented. (e) The change in model evidence by adding specific site variables. Site variables presented in bold increased model evidence and were selected as a covariate (in addition to clinical and demographic covariates). HC = Healthy control; AD = patient using antidepressants; BZ = patients using benzodiazepines; NM = currently unmedicated patients;  ΔISC = difference in inhibitory self-connection strength; DMN = default mode network; SAL = salience network; CEN = central executive network; aPFC = anterior prefrontal cortex; sPar = superior parietal cortex; mPFC = medial prefrontal cortex; dmPFC = dorsomedial prefrontal cortex; iPar = inferior parietal cortex; dACC = dorsal anterior cingulate cortex; PCC = posterior cingulate cortex.

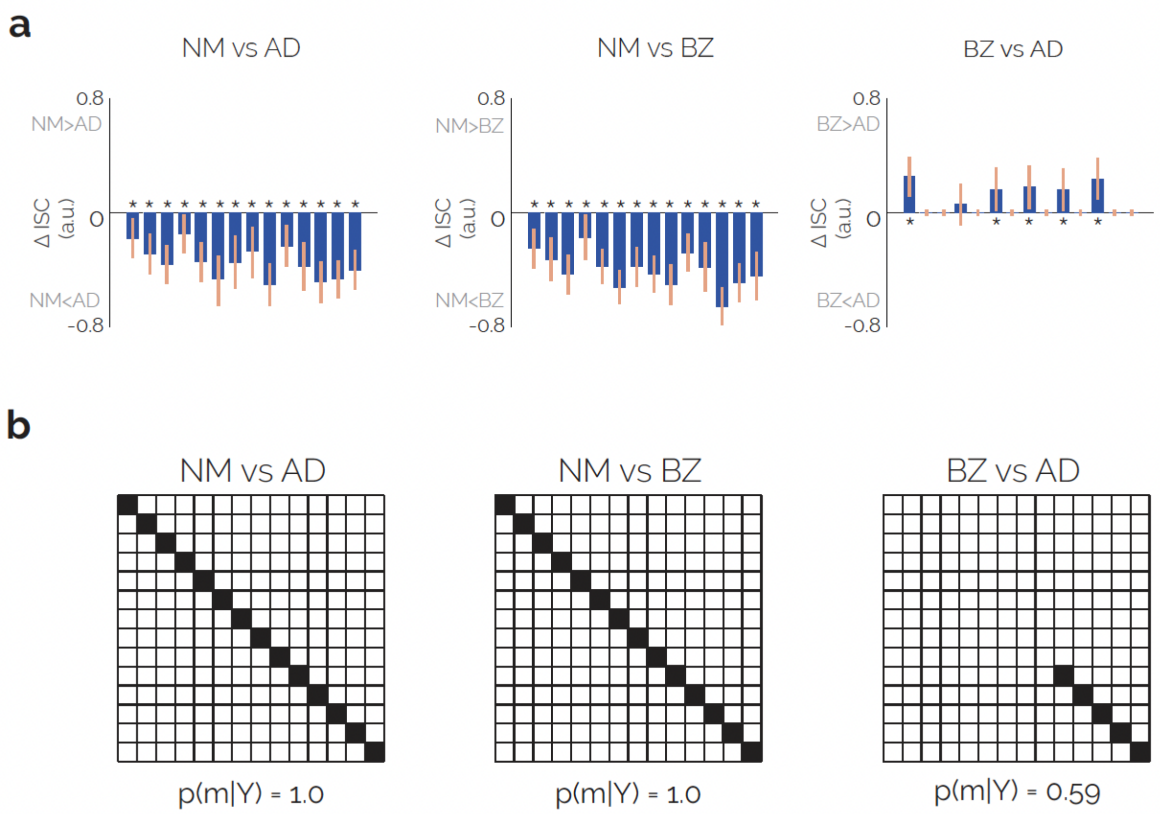


**SI Figure 5 Post-hoc analysis for group differences**

Results of the post-hoc analyses on pairwise comparisons between patient groups. (a) Differences in inhibitory self-connection (ISC) strength of each node between patient groups. Asterisks indicate strong evidence for a group difference, corrected for multiple comparisons. Error bars indicate two standard deviations. The order in which connectivity parameters are presented is identical to the order in the other figures. (b) Winning models for post-hoc comparisons, which indicate strong evidence for a difference in inhibitory self-connections across networks in unmedicated patients compared to patients using benzodiazepines and/or antidepressants. AD = patient using antidepressants; BZ = patients using benzodiazepines; NM = currently unmedicated patients; ΔISC = difference in inhibitory self-connection strength.

SI Fig 6

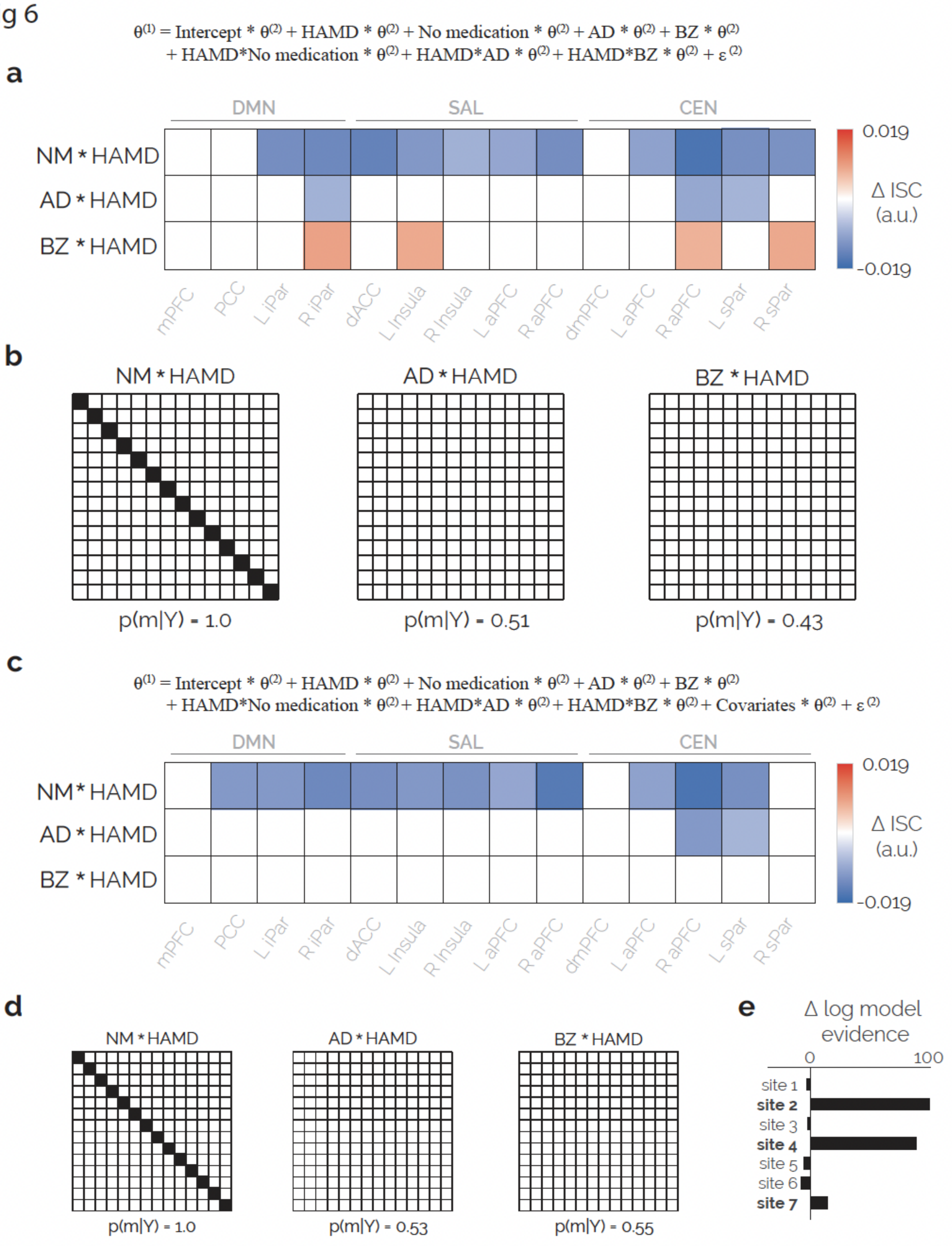


**SI Figure 6 Associations with depression severity**

Results on associations with depression severity are presented. (a) The model design tested for main and interaction effects of depression severity and groups (see SI Figure 3 for further explanation of parameters). Results of automatic search over reduced models (testing for effects in individual connections) is shown. (b) Winning models in the analysis on pre-specified connectivity models is presented for the group by depression severity interaction effects. (c) An additional analysis was performed to test whether results were robust against potential confounders. The model design (identical to (a), plus additional covariates) and the results on the covariate-adjusted automatic search over reduced models are presented. (c) Results of the adjusted analysis on pre-specified models is shown. (e) The change in model evidence by adding specific site variables. Site variables presented in bold increased model evidence and were selected as a covariate (in addition to clinical and demographic covariates). HC = Healthy control; AD = patient using antidepressants; BZ = patients using benzodiazepines; NM = currently unmedicated patients; HAMD = HAM-D score at baseline; ΔISC = difference in inhibitory self-connection strength; DMN = default mode network; SAL = salience network; CEN = central executive network; aPFC = anterior prefrontal cortex; sPar = superior parietal cortex; mPFC = medial prefrontal cortex; dmPFC = dorsomedial prefrontal cortex; iPar = inferior parietal cortex; dACC = dorsal anterior cingulate cortex; PCC = posterior cingulate cortex.

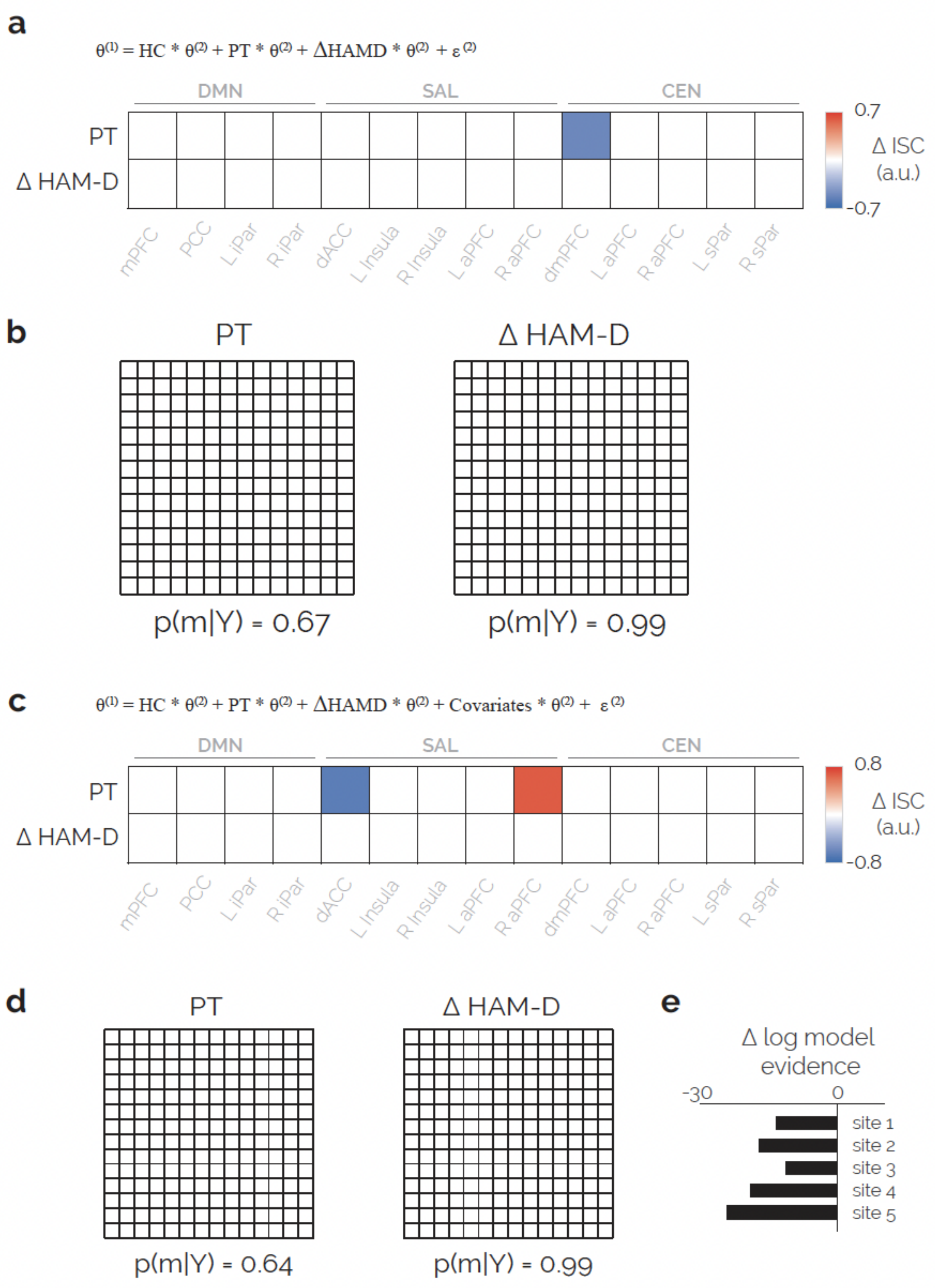


**SI Figure 7 Influence of ECT on inhibitory self-connections and associations with ECT effectiveness**

Longitudinal effects of ECT on self-inhibition and associations with ECT effectiveness were tested in a single model. (a) The model design included healthy controls as the intercept, a regressor for all patients and the change in HAM-D after ECT (compared to baseline). Results of the automatic search over reduced models (testing for effects in individual connections) are shown. The PT regressor models the influence of ECT on inhibitory self-connections, and the ΔMADRS regressor models associations with ECT effectiveness. (b) Winning models in the analysis on pre-specified connectivity models is presented for the effect of patients (i.e., longitudinal effects of ECT, independent from treatment effectiveness) and ECT effectiveness (change in HAM-D). (c) An additional analysis was performed to test whether results were robust against potential confounders. The model design (identical to (a), plus additional covariates) and the results on the covariate-adjusted automatic search over reduced models are presented. (c) Results of the adjusted analysis on pre-specified models are shown. (e) The change in model evidence by adding specific site variables. Site variables presented in bold increased model evidence and were selected as a covariate (in addition to clinical and demographic covariates). HC = Healthy control; PT = all patients; ΔHAMD = change in HAM-D from before to after ECT; ΔISC = difference in inhibitory self-connection strength; DMN = default mode network; SAL = salience network; CEN = central executive network; aPFC = anterior prefrontal cortex; sPar = superior parietal cortex; mPFC = medial prefrontal cortex; dmPFC = dorsomedial prefrontal cortex; iPar = inferior parietal cortex; dACC = dorsal anterior cingulate cortex; PCC = posterior cingulate cortex.

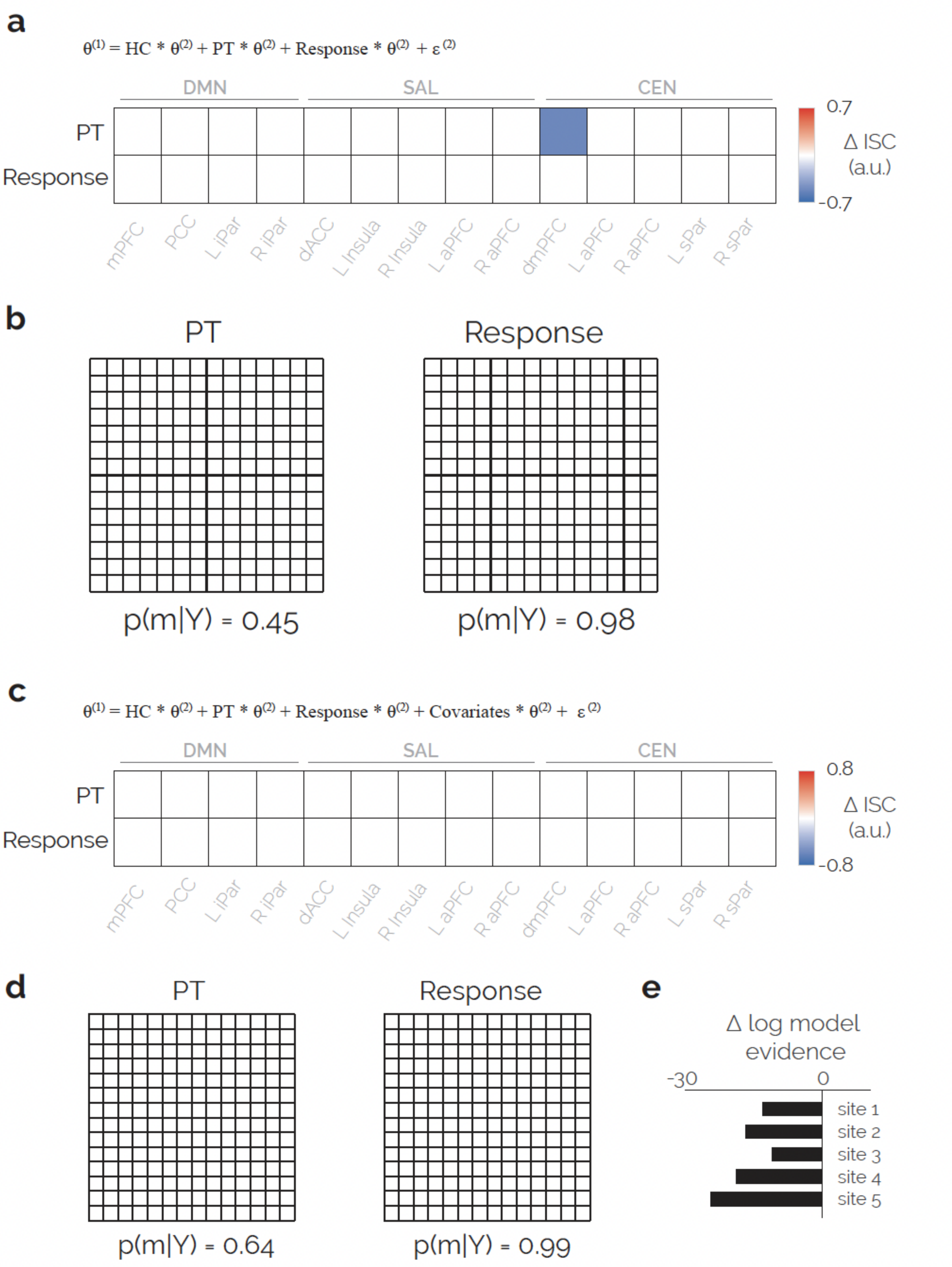


**SI Figure 8 Associations with ECT response**

The analysis is identical to SI Figure 7. However, instead of testing for treatment effectiveness (change in HAM-D) it tests for effects of treatment response (i.e., a binary regressor indicating >50% reduction in HAM-D after ECT). (a) The model design and the results of the automatic search over reduced models (testing for effects in individual connections) are shown. (b) Winning models in the analysis on pre-specified connectivity models are presented for the effect of patients (i.e., longitudinal effects of ECT, independent from treatment effectiveness) and ECT response. (c) An additional analysis was performed to test whether results were robust against potential confounders. The model design (identical to (a), plus additional covariates) and the results on the covariate-adjusted automatic search over reduced models are presented. (c) Results of the adjusted analysis on pre-specified models are shown. (e) The change in model evidence by adding specific site variables. None of the site variables increased model evidence and were therefore not included in the analysis. Clinical and demographic covariates were added as covariates (see SI Methods). HC = Healthy control; PT = all patients; Response = >50% reduction in HAM-D after ECT; ΔISC = difference in inhibitory self-connection strength; DMN = default mode network; SAL = salience network; CEN = central executive network; aPFC = anterior prefrontal cortex; sPar = superior parietal cortex; mPFC = medial prefrontal cortex; dmPFC = dorsomedial prefrontal cortex; iPar = inferior parietal cortex; dACC = dorsal anterior cingulate cortex; PCC = posterior cingulate cortex.

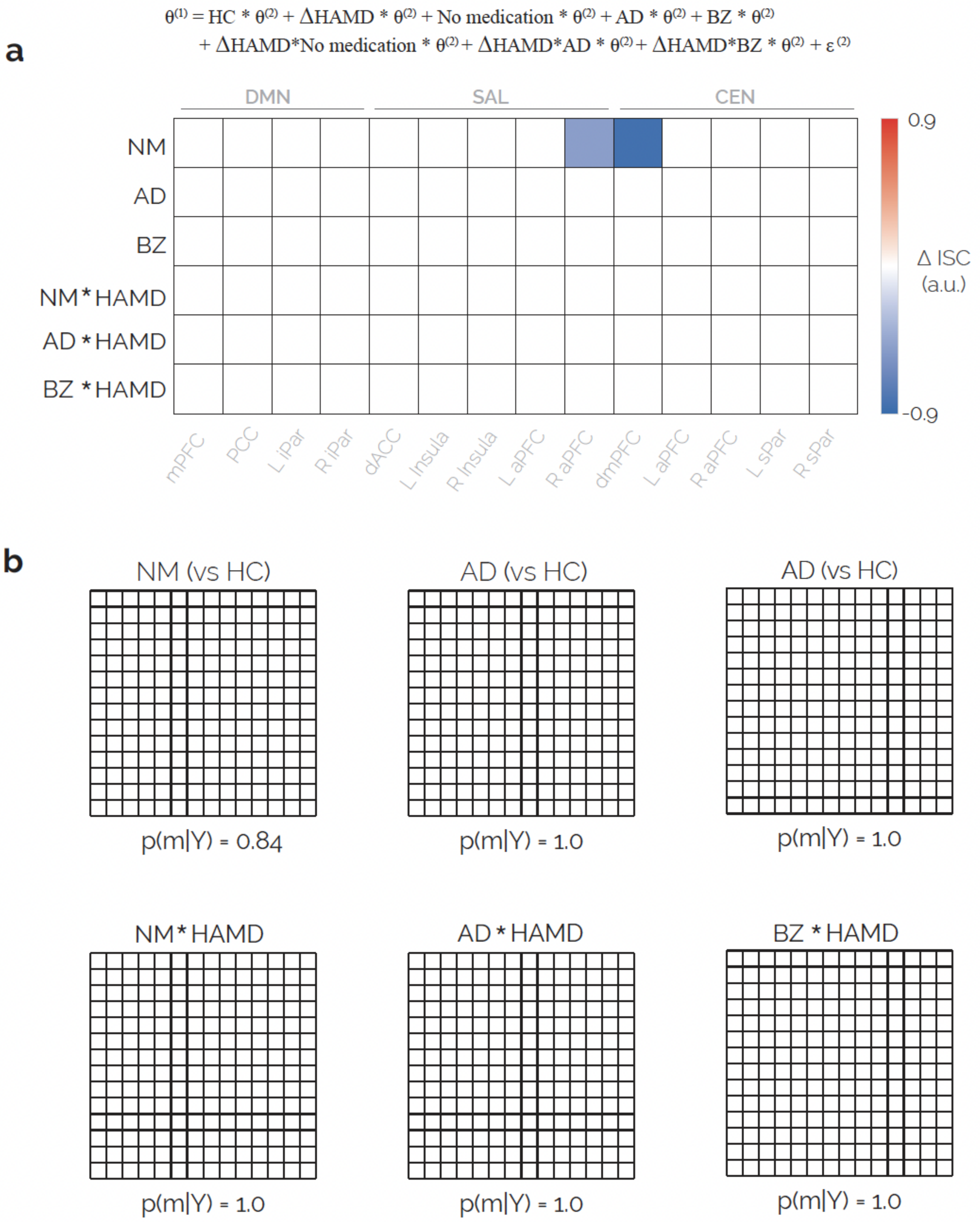


**SI Figure 9 Associations with treatment effectiveness in subgroups**

Additional analysis was run to test for associations with treatment effectiveness in the patient groups based on medication use. (a) The model design included patient group, ECT effectiveness (change in HAM-D), and their interaction effects. The model thereby tested for longitudinal effects of ECT and associations with ECT effectiveness in the distinct patient groups. Effects on individual connectivity parameters are shown. (b) Winning pre-specified connectivity models are shown for longitudinal effects and for associations with treatment effectiveness. HC = Healthy control; AD = patient using antidepressants; BZ = patients using benzodiazepines; NM = currently unmedicated patients; ΔHAMD = change in HAM-D score before and after ECT; ΔISC = difference in inhibitory self-connection strength; DMN = default mode network; SAL = salience network; CEN = central executive network; aPFC = anterior prefrontal cortex; sPar = superior parietal cortex; mPFC = medial prefrontal cortex; dmPFC = dorsomedial prefrontal cortex; iPar = inferior parietal cortex; dACC = dorsal anterior cingulate cortex; PCC = posterior cingulate cortex.
